# Neonicotinoids stimulate H_2_-limited methane emission in *Periplaneta americana* through the regulation of gut bacterium community

**DOI:** 10.1101/2020.09.21.307298

**Authors:** Haibo Bao, Haoli Gao, Jianhua Zhang, Haiyan Lu, Na Yu, Xusheng Shao, Yixi Zhang, Wei Jin, Shuqing Li, Xiaoyong Xu, Jiahua Tian, Zhiping Xu, Zhong Li, Zewen Liu

**Affiliations:** Key laboratory of Integrated Management of Crop Diseases and Pests (Ministry of Education), College of Plant Protection, Nanjing Agricultural University, Weigang 1, Nanjing 210095, China; Institute of Plant Protection, Jiangsu Academy of Agricultural Sciences, Zhongling 50, Nanjing 210014, China; Shanghai Key Laboratory of Chemical Biology, School of Pharmacy, East China University of Science and Technology, Meilong Road 130, Shanghai 200237, China; College of Veterinary Medicine, Nanjing Agricultural University, Weigang 1, Nanjing 210095, China; College of Resources and Environmental Sciences, Nanjing Agricultural University, Weigang 1, Nanjing 210095, China

## Abstract

Methane emitted by insects is considered to be an important source of atmospheric methane. Here we report the stimulation of methane emission in *Periplaneta americana*, an insect species with abundant methanogens, by neonicotinoids, insecticides widely used to control insect pests. The application of cycloxaprid (CYC) and imidacloprid (IMI) caused foregut expansion in *P. americana*, and increased the methane production and emission. Antibiotics could mostly eliminate the stimulatory effects. In *P. americana* gut, hydrogen levels increased and pH values decreased, which could be significantly explained by the gut bacterium community change. The proportion of several bacterium genera increased in guts following CYC treatment, and four genera from five with increased proportions could generate hydrogen at anaerobic conditions. Hydrogen is a central intermediate in methanogenesis. Gut methanogens could use the increased hydrogen to produce more methane, especially at acidic conditions. Following neonicotinoid applications, all increased methanogens in both foregut and hindgut used hydrogen as electron donor to produce methane. Besides, the up-regulation of *mcrA*, encoding the enzyme that catalyzes the final step of methanogenesis, suggested an enhanced methane production ability in present methanogens. In the termite *Coptotermes chaohuensis*, another methanogen-abundant insect species, hydrogen levels in gut and methane emission significantly increased after neonicotinoid treatment, which was similar to the results in *P. americana*. In summary, neonicotinoids changed bacterium community in *P. americana* gut to generate more hydrogen, which then stimulate gut methanogens to produce and emit more methane. The finding raised a new concern over neonicotinoid applications, and might be a potential environmental risk associated with global warming.

## Introduction

Neonicotinoids are extensively used to control insect pests important in both crop protection and animal health. Because of their high efficacy, neonicotinoids have become one of the main classes of insecticides for a range of insect species since the early 1990s. By 2008, neonicotinoids had accounted for 24% of the global insecticide market (Peter et al., 2010) and the market share had increased to more than 25% in 2014 (Bass et al., 2015). However, environmental concerns on neonicotinoids have become increasingly prevalent in recent years, such as the accumulation in soil, leaching into waterways, systemic persistence in crops and plants, and substantial impact on bees and other pollinators (Godfray et al., 2015; Goulson and Kleijn, 2013). For example, neonicotinoids were thought as an important factor for the bee colony collapse disorder due to their lethal and sublethal effects, with reduced learning, foraging, and homing ability (Henry et al., 2012; Stanley et al., 2016), as well as reduced colony growth and production of new queens at the population level (Godfray et al., 2015; Whitehorn et al., 2012). Here, we first reported a new finding on neonicotinoid application stimulating methane production and emission in *Periplaneta americana* (American cockroach), an insect species with abundant methanogens. Because methane emitted by insects is considered to be an important source of atmospheric methane (Yvon-Durocher et al., 2014), the stimulation of methane emission in methanogen-abundant insects by neonicotinoids would be a concern for the environmental risk associated with global warming.

## Results

### Observation of morphological changes of *Periplaneta americana* and determination of H_2_ pressure and pH value in guts

Cycloxaprid (CYC), a new neonicotinoid recently developed and registered in China, shows high insecticidal activity against a range of insect pests (Casida, 2018; Sparks and Nauen, 2015; Zhang et al., 2019). When treated with CYC, *P americana* was stretched and thickened (Fig. 1A), and the foregut was bloated enormously (Fig. 1B). The morphological changes were both time-dependent (Fig. 1C, Fig. S1A) and concentration-dependent (Fig. 1D, Fig. S1B). The morphological changes occurred rapidly after CYC treatment, developed sharply over time and reached the peak at 30 h with an increase of 30% in the body length and 35% in thickness (Fig. S1). Body extension in *P. americana* was also observed in the imidacloprid (IMI) treatment (Fig. 1E), which was also time-dependent and concentration-dependent (Fig. S2).

**Figure 1.**
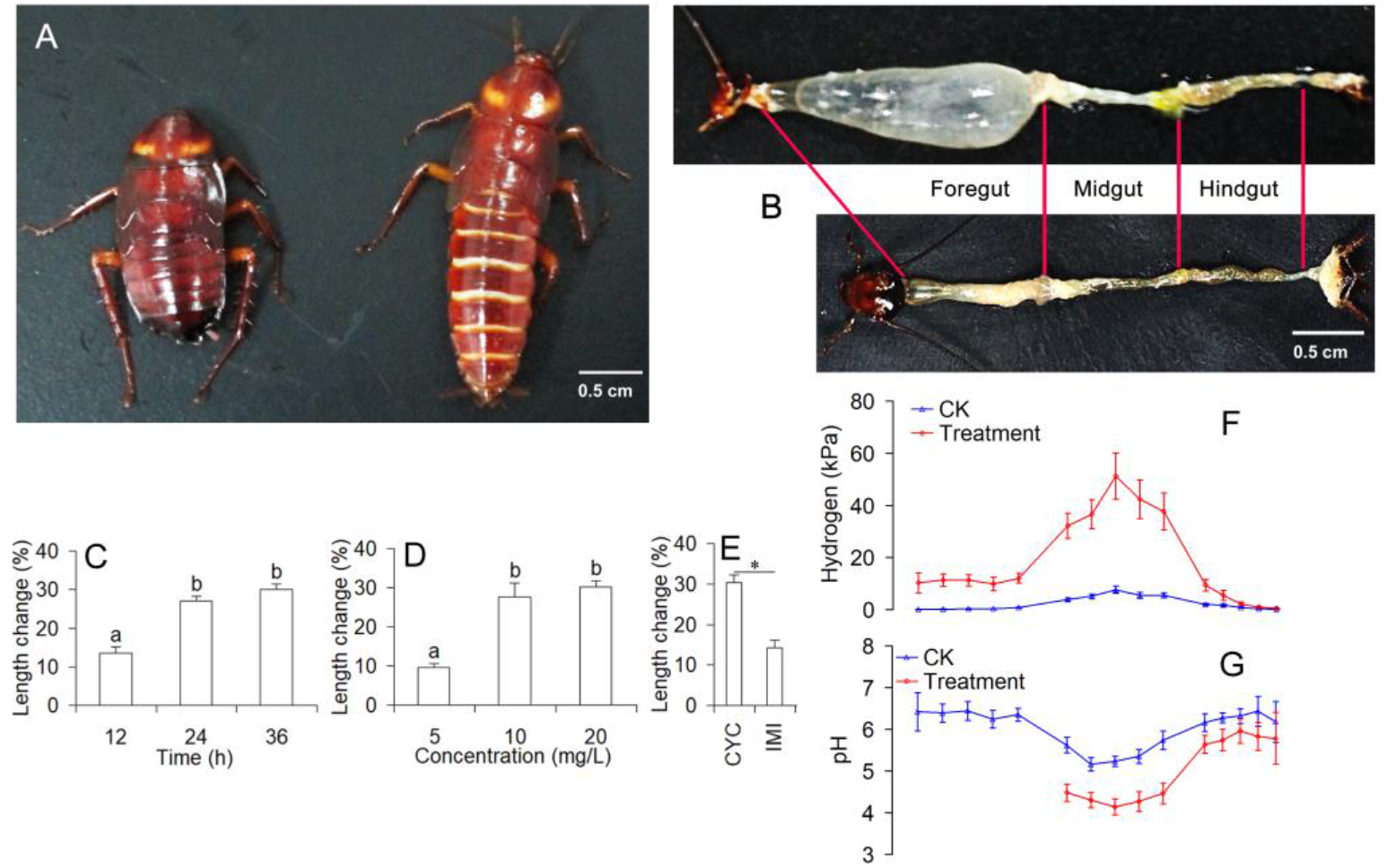
Morphological changes in *Periplaneta americana* following neonicotinoid treatments and changes in gut H_2_ pressure and pH value. (A) The 9th instar nymphs of *P americana* following CYC treatment (right) and the untreated control (left). (B) The dissected guts from CYC-treated (up) and untreated (bottom) *P. americana*. (C) Changes in the body length at different time points after CYC treatment at *LC20* concentration. (D) Body length changes at 36 h after the CYC treatment at different concentrations. In C and D, different letters indicated significant differences at 0.05 level. (E) Body length changes at 36 h following CYC and IMI treatments at *LC*_20_ concentrations. *, significant difference at 0.05 level. (F) H_2_ pressure comparison between CK and CYC treatment. (G) pH value comparison between CK and CYC treatment. In F and G, the abscissa of each test site in insect gut corresponded to that of CK gut in B. Because pH micro-sensor tip could not position to the measure site in the bloated foregut, pH data of foregut were not obtained in CYC treated insects (G). In C-G, data are mean±SEM from at least five repetitions.

H_2_ pressure and pH value were determined in CYC treated *P. americana* gut and compared to that of untreated control. In the untreated *P. americana* gut, H_2_ pressure was close to 0 (Fig. 1F). The greatest increase of H_2_ pressure was 51 kPa in the midgut center following CYC treatment. However, the increase ratios in foreguts (14.8-85.6 times) were much bigger than that in midguts (6.8-8.4 times) and hindguts (2.2-4.7 times) when compared to CK (Fig. 1F). The pH values in CYC treated *P. americana* midguts and hindguts significantly decreased when compared to untreated control (Fig. 1G). The pH data were not successfully determined in foreguts of CYC-treated insects, due to the difficulty in positioning micro-sensor tip to the measure site in the bloated gut (Fig. 1G). The bloated gut burst when the pH micro-sensor tip penetrated the up site and moved to the bottom site where the pH measure content was.

### Stimulation of methane emission from *P americana* by neonicotinoids

The gas in bloated foreguts of *P. americana* was collected and determined by gas chromatography. A single peak was observed at the retention time of 1.873 min (Fig. 2A), which was almost identical to that of methane standard (Fig. S3A). Then methane emission was determined and quantified. CYC and IMI at *LC*_20_ stimulated methane emission in a time-dependent manner, and the emission rate reached the peak (37.3 ng/insect/h) at 30 h for CYC and (23.3 ng/insect/h) at 33 h for IMI, which was 17.4 and 6.33 times as much as the untreated control, respectively (Fig. 2B). Concentration-dependent stimulation of the methane emission was observed for both CYC and IMI (Fig. 2C). The antibiotics chloramphenicol (Cp) and N-[2-(Nitrooxy) ethyl]-3-pyridinecarboxamide (NPD) could mostly eliminate the stimulatory effects on methane emission by insecticide treatments (Fig. 2D).

**Figure 2.**
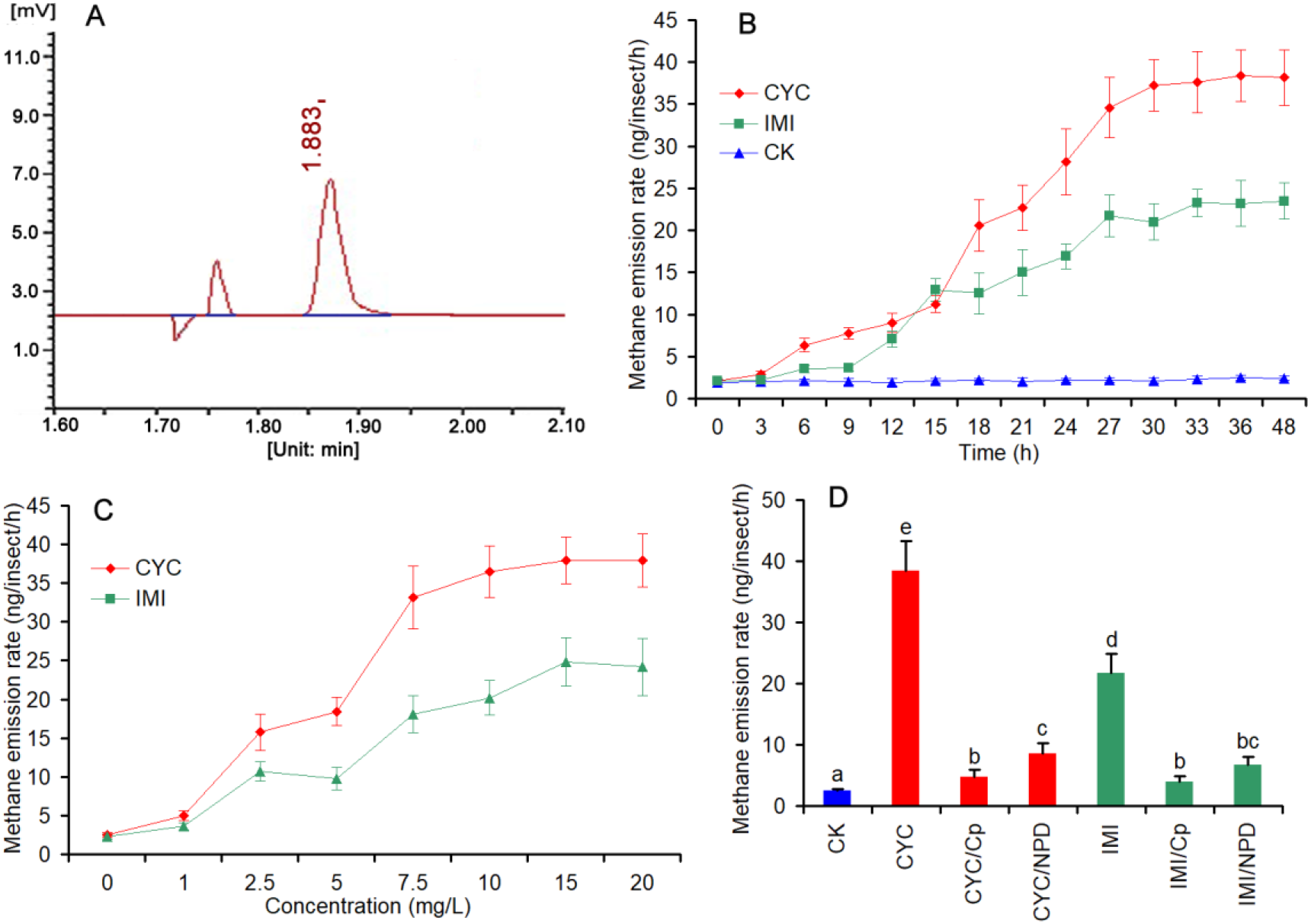
Identification and quantitative determination of methane from *P. americana* following neonicotinoid applications. (A) The gas chromatography identification of methane in the bloated foreguts of *P americana* following CYC treatment. (B) Methane emission from *P americana* at different time points following CYC and IMI application at *LC*_20_ and untreated control. (C) Methane emission from *P. americana* at 36 h following CYC and IMI treatment with different concentrations. (D) Effects of antibiotics on methane emission from *P. americana* following the treatment by CYC and IMI at *LC*_20_ concentration. To calculate the emission quantity of methane, the relationship line between chromatography peak area and methane quantity was set up using standard methane (Fig. S3B). In B, C and D, data are mean±SEM from at least five repetitions.

### Stimulation of methane emission from *Coptotermes chaohuensis* by neonicotinoids

The methane emission was also stimulated by neonicotinoids in the wingless workers of *C*. *chaohuensis* (Figure 3A), although neither morphological change nor gut bloat was observed (Fig. 3B, Fig. S4). At *LC*_20_ concentration, CYC had higher stimulatory effects on methane emission than that of IMI, which was similar to that in *P. americana*. H_2_ pressure was also determined in CYC treated *C*. *chaohuensis* gut. Similar to results of *P. americana*, H_2_ pressure significantly increased in CYC-treated *C*. *chaohuensis* gut when compared to untreated control (Fig. 3C). However, the increase in H_2_ pressure was negligible at the end of hindgut.

**Figure 3.**
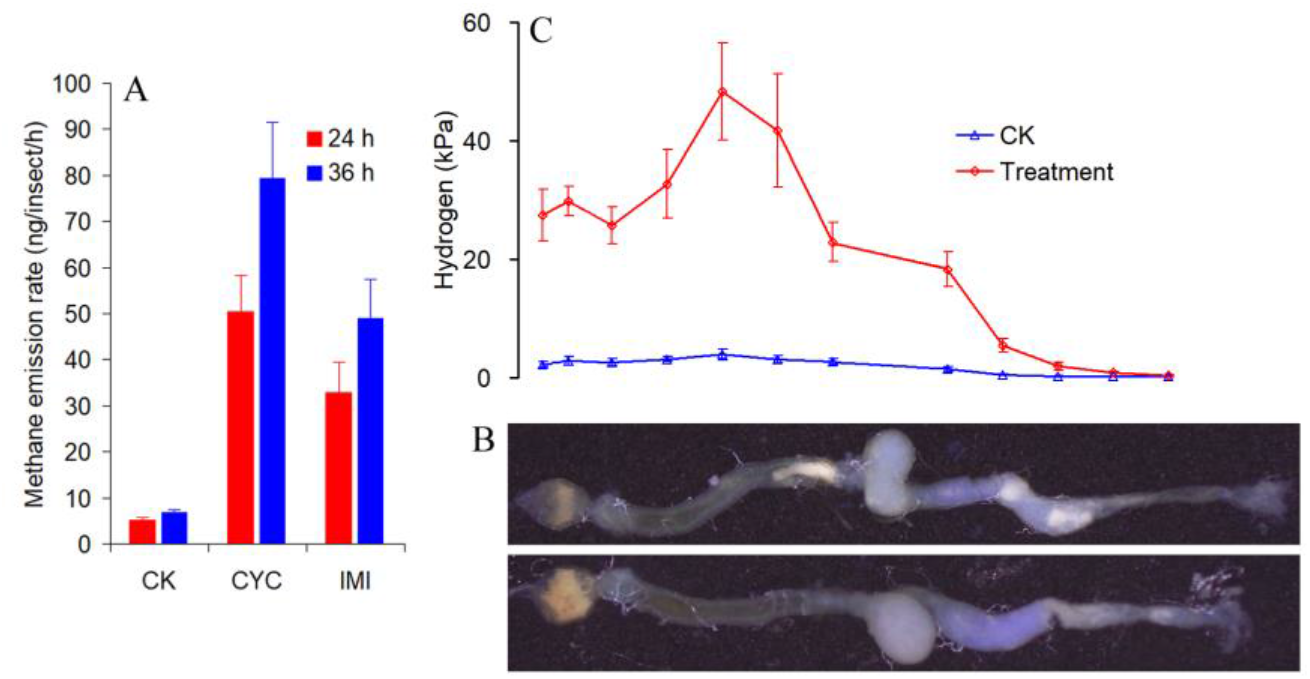
Determination of hydrogen pressure in gut and methane emission of *C*. *chaohuensis*. (A) Methane emission from *C*. *chaohuensis* at 24 h and 36 h after CYC and IMI application at *LC*_20_ concentration and untreated control. (B) The dissected guts from CYC-treated (up) and untreated (bottom) *C*. *chaohuensis*. (C) Gut H_2_ pressure comparison between CK and CYC treatment. In C, the abscissa of each test site in insect gut corresponded to that of gut present in B. In A and C, data are mean±SEM from at least five repetitions.

### Effects of neonicotinoids on methane emission from soils and sheep rumen fluids

Due to fail in culturing methanogens from *P. americana*, it was impossible for us to determine whether *P. americana* gut methanogens was in direct response to neonicotinoid treatment. We instead collected methanogens from the rumen fluid of the Chinese Hu sheep for insecticide treatments^15^. CYC treatment on rumen fluids neither changed the total amount of gas emission (Fig. 4A) nor increased methane emission (Fig. 4B). CYC at the concentration up to 500 mg/L reduced methane emission was possibly due to the inhibition on methanogen population. The results showed that neonicotinoids did not stimulate methane production in the collected methanogens from rumen animals. However, whether neonicotinoids change methane production in the rumina *in vivo* needs further studies.

**Figure 4.**
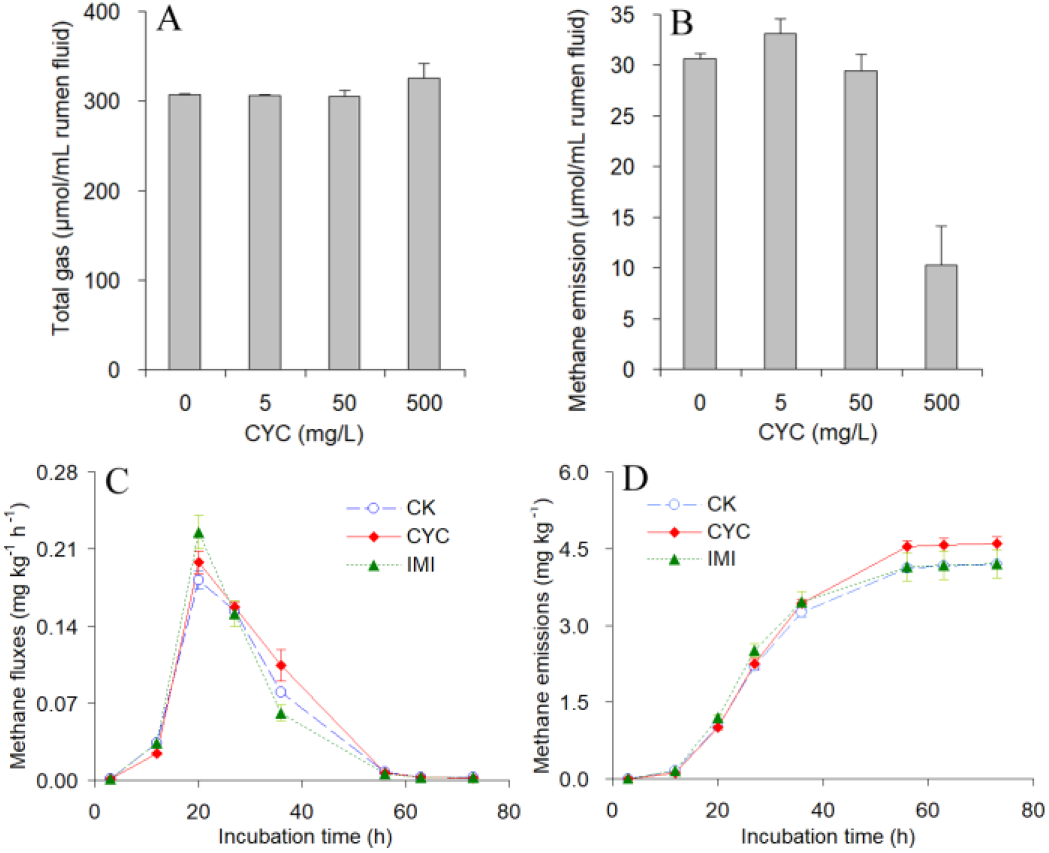
Effects of neonicotinoids on methane emission from rumen fluids of the Chinese Hu sheep and rice field soils. Total gas (A) and methane (B) emission mounts were determined in sheep rumen fluids treated by different concentrations of CYC. Methane emission rate (C) and mount (D) were determined in rice field soils with addition of CYC and IMI at a final concentration of 500 mg/kg soil. Data are mean±SEM from at least three repetitions.

Neonicotinoids can persist and accumulate in soil, and rice fields are major anthropogenic sources of methane emission^3, 16^. If neonicotinoids stimulate methane production in methanogens in rice field soil, it may have an impact on the environment. Rice field soil was collected and treated with CYC and IMI^17^. Fortunately, no significant change in the methane production and emission from the rice field soil with methanogens was observed when the methanogens were directly treated by either cycloxaprid or imidacloprid (Fig. 4C, 4D).

### Analysis of the bacterium community in *P. americana* guts

The diversity changes of bacteria were estimated using 16S rRNA sequencing in three parts of *P. americana* guts, the foregut, midgut, and hindgut, and compared between CYC-treated individuals and the untreated control (Fig. S5). In three gut parts, CYC treatment caused significant changes in some bacterium genera (Fig. 5). For examples, *Shimwellia* increased from 17.56% to 45.93% in foreguts (Fig. 5A), *Lactococcus* from 0.16% to 10.68%, *Pediococcus* from 0.81% to 8.75% and *Shimwellia* from 0.02% to 7.03% in midguts (Fig. 5B), and *Desulfovibrio* from 5.90% to 12.02% in hindguts (Fig. 5C). Similarly, some bacterium genera in guts decreased after CYC treatment, such as the genus *Arthromitus* (g_Candidatus_*Arthromitus*) from 34.04% to 2.61% in hindguts (Fig. 5C).

**Figure 5.**
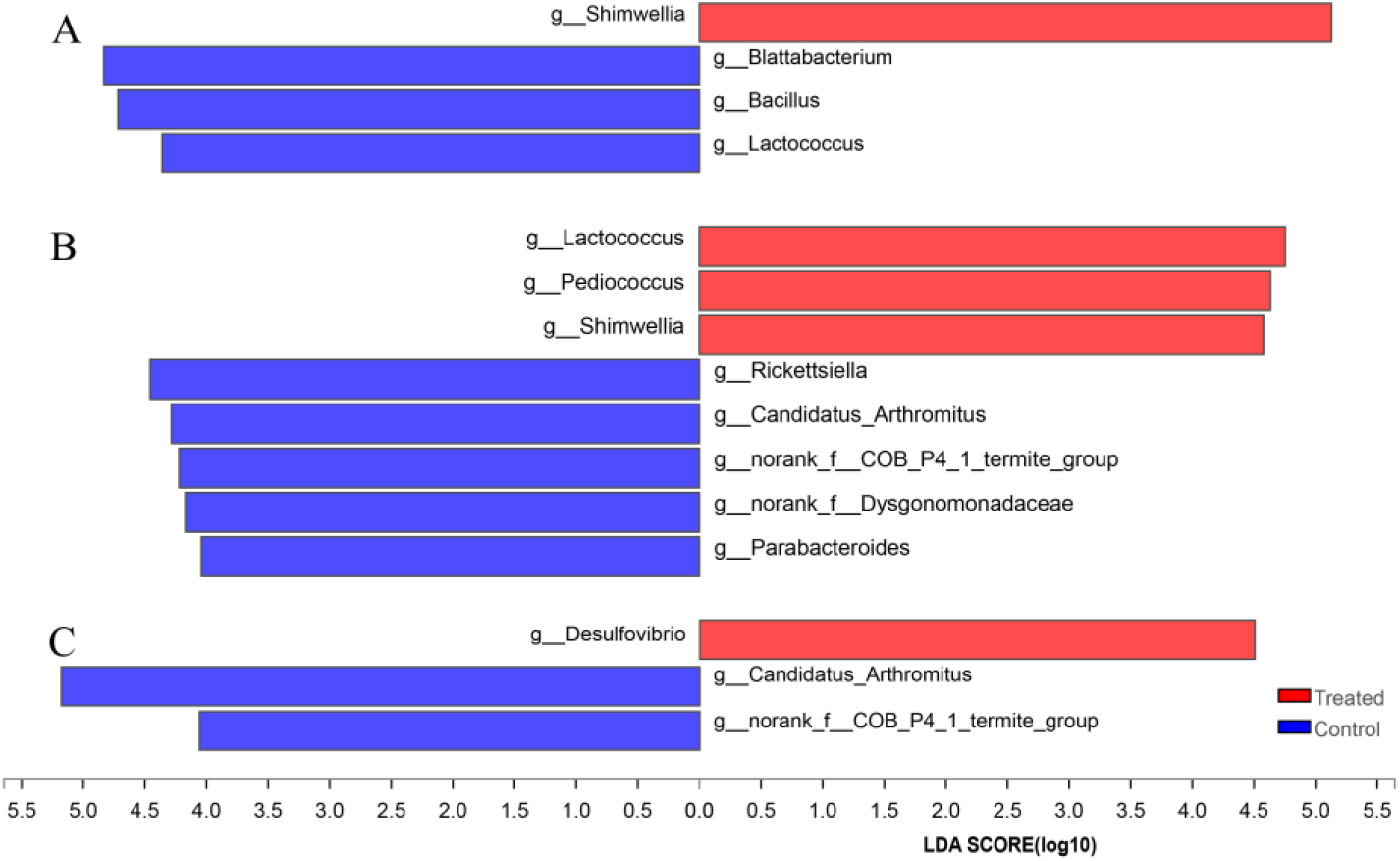
Comparison of bacteria community in three parts of *P. americana* guts between CYC treatment and untreated control. The most different taxa in abundance in bacteria between groups were determined by LDA effect size (LEfSe) analysis, lowercase letters before taxon name mean different assigned taxa, i.e. g for genus and f for family. The threshold of LDA score is 4. (A) Foregut. (B) Midgut. (C) Hindgut.

### Analysis of the archaea and methanogen community in *P. americana* guts

The archaea and methanogen communities were estimated and compared between CYC treatment and untreated control (Fig. S6). Although several trials were performed for midgut samples from both treatments and controls, we could not get enough data to estimate archaea and methanogen communities in this gut part.

In *P. americana* foregut, CYC treatment increased the proportion of the identified methanogens in archaea community when compared to that of untreated control, mainly reflecting in the significant increase of *Methanocorpusculum* (Fig. 6A), the obvious but not significant increase in *Methanobrevibacter*, unclassified_f_*Methanobacteriaceae* and the decrease in unclassified_k_norank_d_*Archaea* (Fig. S6A). The methanogen community of foregut did not significantly change (Fig. S6B), although *Methanobacteria* and *Methanobacteriales* (also belonging to class of Methanobacteria) had significant changes (Fig. 6B), however, with negligible proportions in both CYC treatment and untreated control (Fig. S6B).

**Figure 6.**
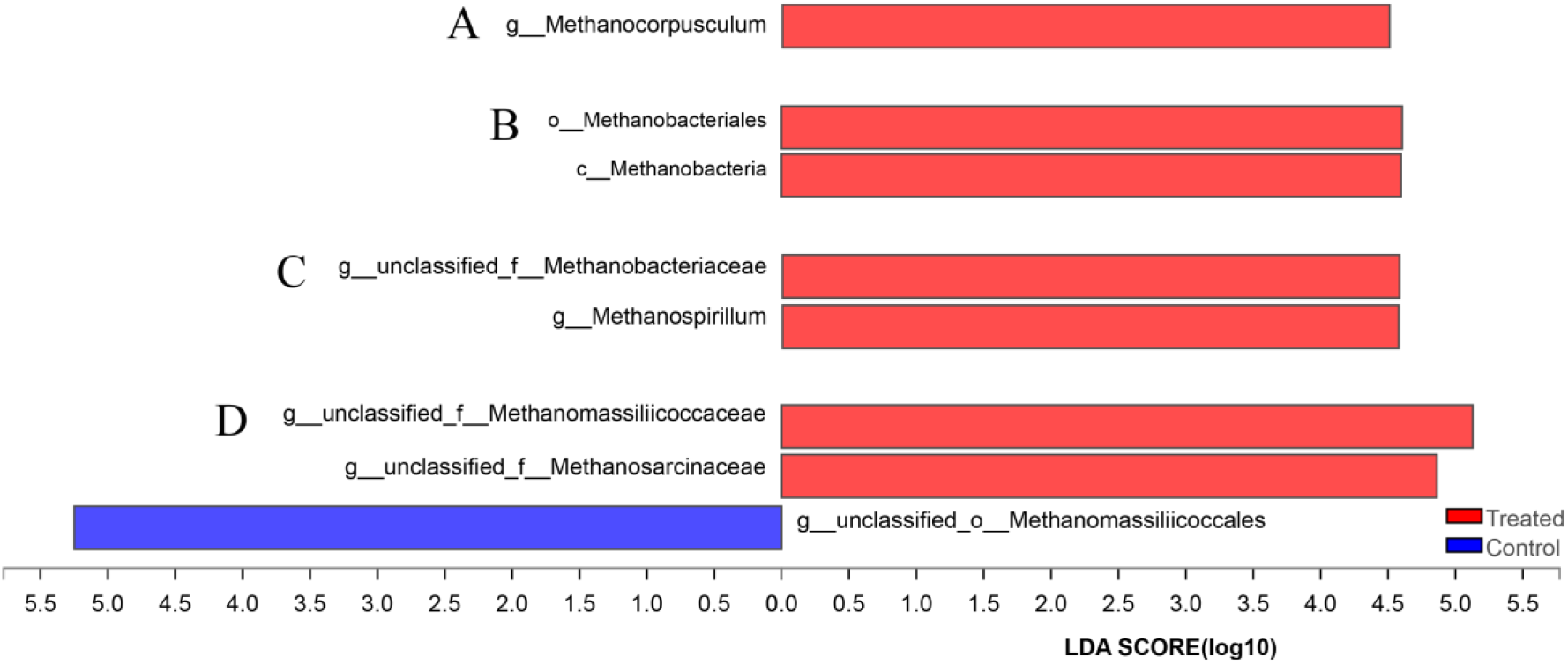
Comparison of archaea and methanogen communities in *P. americana* foreguts and hindguts between CYC treatment and untreated control. The most different taxa in abundance in archaea and methanogens between groups were determined by LDA effect size (LEfSe) analysis, lowercase letters before taxon name mean different assigned taxa, i.e. g for genus, f for family, o for order and c for class. The threshold of LDA score is 4. (A) Archaea in foregut. (B) Methanogens in foregut. (C) Archaea in hindgut. (D) Methanogens in hindgut.

In hindguts, methanogens almost occupied 100% of the archaea community (Fig. S6C). The relevant abundance of two taxa, genus of *Methanospirillum* and family of *Methanobacteriaceae* (no genus classification), significantly increased (Fig. 6C). The methanogen community in hindguts had various changes following CYC treatment (Fig. S6D), with significant increases in *Methanomassiliicoccaceae* and *Methanosarcinaceae*, and decrease in *Methanomassiliicoccales* (Fig. 6D).

### Quantitative analysis of *mcrA* in insect guts following neonicotinoid treatments

The transcript level of *mcrA* was quantified in *P. americana* guts and compared between neonicotinoid treatments and untreated controls. The *mcrA* transcript levels remained stable in wild *P. americana* guts, but increased in insecticide treatments in a time-dependent manner and reached the peak at 36 h (Fig. 7A). CYC and IMI treatments increase *mcrA* transcript levels in guts to 8.01- and 3.97-fold than that of untreated control at 36 h, respectively. The *mcrA* level increase caused by insecticide treatments occurred in the hindgut and foregut, but not in the midgut (Fig. 7B). CYC treatment resulted in an 8.97- and 6.42-fold increase in *mcrA* levels in hindguts and foreguts, and the fold change for IMI treatment were 3.70 and 3.78, respectively.

**Figure 7.**
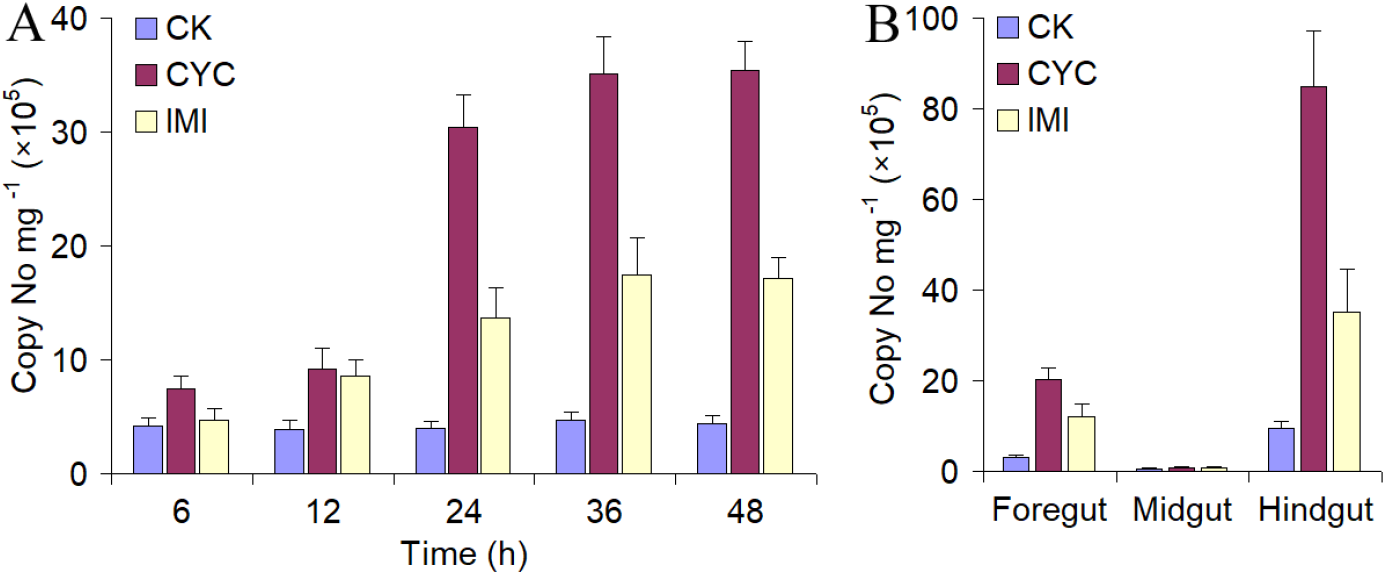
Determination of *mcrA* abundance in *P. americana* guts following CYC and IMI treatments. (A) Changes of *mcrA* abundance in *P. americana* guts at different time points after insecticide treatments. (B) Changes of *mcrA* abundance in different gut parts at 36 h after insecticide treatments. Data are mean±SEM from at least five repetitions.

## Discussion

Insecticides help us prevent crop lose from insect pests, but also bring a dozens of environmental concerns. Neonicotinoids accounted for over one-quarter of the global insecticide market (Bass et al., 2015). However, neonicotinoids were also considered a main factor for the bee colony collapse disorder (Henry et al., 2012; Whitehorn et al., 2012). The methanogen-abundant insects, such as termites, cockroaches, and scarab beetles, emit methane produced by their gut methanogens. Methane emitted by insects is an important source of atmospheric methane (Brune, 2010). Take termites as a representative example, they contribute approximately 3% to global methane emission (Sanderson, 1996). Although these insects are not target pests of neonicotinoids, they are often exposed to neonicotinoids due to neonicotinoid accumulation in soil, leaching into waterways and systemic persistence in plants (Goulson and Kleijn, 2013). Neonicotinoids of the exposure concentrations may be sublethal, but still have the possibility to cause some effects on these insect species, as their sublethal effects on bees (Henry et al., 2012). Here we reported a finding that neonicotinoids, at their sublethal concentrations, stimulated methane emission in the American cockroach, *P. americana*, a methanogen-abundant insect species. The stimulation on methane emission by neonicotinoids was also observed in the termite *Coptotermes chaohuensis*, another methanogen-abundant insect species. The stimulation of methane production and emission was not a general methanogen response to neonicotinoid application, because neonicotinoids treatments did not cause any changes in methane emission in rumen fluids of the Chinese Hu sheep and rice field soils with abundant methanogens. The results revealed that the stimulation of methane production in *P. americana* gut through an indirect way.

Two neonicotinoids, CYC and IMI at *LC*_20_ concentration, changed the bacterium community in *P. americana* guts directly or indirectly, and some of the increased bacterium genera could produce more hydrogen and change the pH condition in gut. *Shimwellia*, a genus with significantly increased proportion in CYC-treated *P. americana* foreguts and midguts, belongs to the family *Enterobacteriaceae* which produces typical acidic fermentation products such as hydrogen and lacatate (Bauer et al., 2015). *Lactococcus*, a genus produces hydrogen at anaerobic conditions (Pandey et al., 2019), had increased proportion in CYC-treated midguts, but was with lower proportion in the treated foreguts. *Desulfovibrio*, a genus with significantly increased proportion in CYC-treated hindguts, produced hydrogen, and they also have the ability to live in stable syntrophic associations with H2-scavenging methanogenic partners (Baffert et al., 2019). *Pediococcus*, with increased percentage in CYC-treated midguts, could lead to pH decrease by producing organic acids, although they did not produce hydrogen (Banwo et al., 2013).

Hydrogen is a central intermediate in methanogenesis and the most important electron donor for hydrogenotrophic methanogenesis (Brune, 2010). In CYC-treated *P. americana* gut, methanogens had two ways of taking full advantage of increased hydrogen, changing methanogen community and increasing methanogenesis ability in the present methanogens. In the hindguts of *P. americana*, which do not accumulate hydrogen, methanogenesis is severely hydrogen-limited (Lemke et al., 2001). Several methanogen genera had higher proportions in CYC-treated *P. americana* guts (foreguts and hindguts) than that in untreated control, and all these methanogen genera belongs to hydrogenotrophic methanogenesis using hydrogen as the electron donor to generate methane (Kroeninger et al., 2019; Thauer et al., 2008). Methanogens are phylogenetically diverse, but the enzyme mcrA is considered typical of all known methanogens (Friedrich, 2005). In some samples, such as the anaerobic digesters treating sludge and wastewater samples, the *mcrA* gene could be recognized as a biomarker for methane yield (Wilkins et al., 2015). In the present study, *mcrA* was significantly up-regulated by both CYC and IMI in *P. americana* guts (foreguts and hindguts), which indicated that methanogen community in CYC-treated guts had a higher ability in methanogenesis than that of untreated control.

Methanogens are generally considered to be restricted in the enlarged hindgut compartment fueled by hydrogen (Brune, 2010). The accumulated hydrogen in guts is toxic to insects, and insects from non-methane-emitting taxa emit substantial amounts of hydrogen and some individuals from methane-emitting taxa that fail to produce methane often emit hydrogen instead (Hackstein and Stumm, 1994; Schmitt-Wagner and Brune, 1999; Sugimoto et al., 1998). Most methanogens have not been cultured yet, and their diversity could only be deduced from the analysis of their 16S rRNA genes (Hackstein and van Alen, 2018). By 16S rRNA sequencing, the diverse methanogens were estimated in both *P. americana* foreguts and hindguts, not restricted in hindgut compartment. *P. americana* does not have an enlarged hindgut compartment, and its foregut has a comparable size to the hindgut. This is the main reason that this gut part was denoted as foregut, but not crop as in most methanogen-abundant insects (Brune, 2010). *P. americana* foregut and hindgut have similar hydrogen pressure, which is much lower than that of its midgut. The distinct gut structure of *P. americana* make it possible to contain methanogens in the foregut. Successful detection and quantification of *mcrA* gene in foreguts also provided indirect evidences for the presence of methanogens in *P. americana* foreguts (Friedrich, 2005). In future, we will try to find out other direct evidence to confirm the presence of methanogens in *P. americana* foreguts.

An interesting finding in this study was the bloated foregut in *P. americana* after neonicotinoid treatment. Methane was detected in the bloated foregut gas, which provided a direct clue for us to test methane production and emission from *P. americana* following insecticide applications. A possible explanation for the bloated foregut is that methane was massively produced by foregut methanogens using hydrogen in foregut itself and from midgut. As results shown, the hydrogen pressure in neonicotinoid treated midgut and foregut was significantly higher than that in untreated control. The *P. americana* foregut gas could not be emitted from its esophagus and cross midgut, and then caused the foregut bloat. *P. americana* hindgut does not accumulate hydrogen as well as methane (Lemke et al., 2001), so the bloat appearance was not observed in its hindgut. In *P. americana* hindgut, methane was generated by methanogens using hydrogen and emitted by its anus.

At anaerobic conditions in insect guts, the accumulated hydrogen inhibits the degradation of organic matters by microbes, due to the difficulty in cyclical utilization of NAD (nicotinamide adenine dinucleotide) and the abnormity in oxidation-reduction reaction. So the accumulated hydrogen in guts affected adversely on insects, although the effects may be sublethal and without direct toxicity such as quick death (Hackstein and Stumm, 1994). In methanogen-abundant insects, methanogens effectively utilize the generated hydrogen, which is an important mechanism to detoxify insects from accumulated hydrogen in guts. If neonicotinoids also stimulate hydrogen generation in insects without methanogens in guts, such as in bees, is it possible that the high hydrogen concentration leads to sublethal effects on insect physiology and population growth?

## Materials and methods

### Insects and Chemicals

*Periplaneta americana* was purchased from Feitian Medicinal Animal Co. Ltd. (Danyang, China). *Coptotermes chaohuensis* was provided by Nanjing Termite Control Institute (Nanjing, China). Cycloxaprid (CYC) was provided by East China University of Science and Technology (Shanghai, China). Imidacloprid (IMI) was purchased from Sigma-Aldrich (St. Louis, MO). Chloramphenicol (Cp) was purchased from ABC One (Shanghai, China). N-[2-(Nitrooxy) ethyl]-3-pyridinecarboxamide (NPD) was obtained from Nipro Pharma Corporation Kagamiishi Plant (Kagamiishi, Japan).

### Bioassay

The bioassays were performed with the oral application method. Insecticides were dissolved in acetone and then diluted to required concentrations with distilled water. Ten microliter insecticide solution was orally applied to the 9th instar nymphs of *P. americana* with a 10 μL pipette. *C*. *chaohuensis* (wingless worker) were fed with insecticide solutions soaked in a piece of filter paper for 12 h. Aqueous acetone solution equivalent to the maximum volume in all insecticide solutions was used as the control. The treated insects were maintained in the greenhouse and the mortality was recorded at 48 h. For the antibiotic treatment, Cp and NPD were applied at the concentration of 5 mg/L orally applied to *P. americana*. Insecticides were then applied at 72 h after the antibiotic treatments.

### Recording changes in *P. americana* bodies and guts

The lethal concentration 20% (*LC*_20_) of each insecticide on *P. americana* was calculated based on the bioassay results. Insecticides at the concentration of *LC20* were applied in further experiments. The insect body length and thickness were then measured with a vernier caliper at a serial of time points. The intact guts were dissected and photographed at 24 h after the insecticide treatment. At least fifteen insects were measured with.

### Detection and quantitative analysis of methane

To detect gas in the foregut of *P. Americana* after neonicotinoid applications, five intact foreguts were dissected and deposited in a 5 mL airtight vial. The vial was fully filled with pure nitrogen, and then sealed with a rubber plug. The foregut gas was released with a 2.5 mL gas-tight syringe and fully mixed. Then 1 mL gas was collected and rapidly injected into the chromatographic system and recorded on Agilent 6820 with chromatographic column Agilent HP-AL/S (30m×0.530mm×15nm). Injection conditions were at the column temperature 80 °C, vaporization chamber temperature 100 °C, hydrogen ion flame detector (FID), detection temperature 230 °C, carrier gas nitrogen, column flow rate 2.9 mL/min, and split ratio 10:1.

To collect the methane emitted by insects, the test insects were incubated in a 500 mL glass bottle. The bottle had a metal lid with an opening (10 mm diameter) plugged with a butadiene rubber septum. Thirty 9th instar nymphs of *P. Americana* or 150 wingless workers of *C*. *chaohuensis* were used in each bottle. The gas samples (2.5 ml) were aspirated by a syringe through the septum for the methane concentration determination at different time points following the incubation. Using the methane standard, the calculation equation between the methane mass and areas in chromatographic recording was determined as y=0.0014x+1.6663, in which y is the methane quantity (ng) and x is the peak area (uV×s).

### Determination of H_2_ pressure and pH values in *P. americana* gut

The intestinal H_2_ pressure was measured according to Hydrogen Sensor User Manual using the standard glass micro-sensor H2-50 (outside tip diameter 40 μm, Unisense, Aarhus, Denmark). The micro-sensor was fixed on a micromanipulator and calibrated in Ringer’s solution containing a 95 % N_2_ and 5 % H_2_. The pH value was determined according to pH and Reference Electrode Manual using the standard glass micro-sensor pH-50 (outside tip diameter 50 μm, Unisense, Aarhus, Denmark). The pH microelectrode was calibrated with the commercial pH standard solutions of pH 3.0, 5.0, 7.0 and 9.0. The dissected *P. americana* and *C. chaohuensis* guts were placed in a small PVC chamber and was irrigated with air-saturated Ringer solution at pH 7.0. The microelectrode was moved with a minimum step increment of 50 μm and the tip was positioned with a horizontally mounted stereomicroscope. All measurements were carried out at 25 °C. For each part of the gut (foregut, midgut, and hindgut), five sites were selected for the measurement. The test at each site was repeated for at least 5 times.

### Determination of effects of neonicotinoids on rumen fluids of Chinese Hu sheep

Three rumen-fistulated Chinese Hu sheep were used as inoculant donor animals. The experiment was performed according to the procedure reported by Martínez-Fernández et al. (Martínez-Fernández et al., 2014). Alfalfa was milled to 1 mm long before being weighed into 100 mL serum bottles. Ruminal contents were obtained immediately before the morning feeding from the three sheep, pooled, and strained through 4 layers of cheesecloth into an insulated flask under anaerobic conditions. The filtered rumen fluid was mixed with the buffer solution at a ratio of 1:3 (v:v) at 39 °C under anaerobic conditions(Menke and Steingass, 1987). Each bottle contained 0.5 g alfalfa and 50 mL buffered rumen fluid. Chemicals were first dissolved in DMSO and then directly added into the bottles before the inoculation with the final DMSO concentration less than 0.1%. Bottles were sealed with rubber stoppers and aluminum caps and incubated at 39 °C for 24 h. Each set of experimental tests had four replicates. Gas production was measured using a pressure transducer and a calibrated syringe (Theodorou et al., 1994). Following the gas measurement, methane and hydrogen production was determined immediately with a GC-TCD instrument (Agilent 7890B, Agilent, California, USA) according to the method described by Jin et al. (Jin et al., 2017).

### Determination of effects of neonicotinoids on rice field soils

Soil from the plow layer (0-20cm) was collected from a rice field in Jurong (China) in December, 2017. Soil was dried, sieved (2-mm mesh size), mixed and stored at room temperature. In a 25-mL glass vial, 2 g of dried soil was placed and flushed with N_2_ for 20 min to create an anaerobic environment. The insecticide was first dissolved in DMSO and applied to the soil in vials at a final concentration of 500 mg/kg soil. Four to five replicates were conducted in each experiment. The sample bottles were incubated in the dark at 30°C for 7 days (Zou et al., 2004). To determine CH_4_ emissions, each incubation vial was ventilated by flushing the headspace with N_2_ for 20 min and then sealed to accumulate CH_4_. After subsequent anaerobic incubation of 6-8 h, gas samples were obtained from the headspace of incubation vials using miniaturized gas samples to determine CH_4_ emission rates. CH_4_ concentrations were measured within 24h using a gas chromatograph (Agilent 7890A) coupled with thermal conductivity detector and flame ionization detector. The oven was operated at 55 °C, the FID at 200 °C(Zou et al., 2004).

### Analysis of microbial community diversity

The foreguts, midguts, and hindguts of 40 *P. americana* individuals treated with CYC at *LC*_20_ (calculated from the toxicity bioassay results, Table S1) or untreated control (CK) were grouped into one sample, respectively. Three independent samples for each treatment and control were prepared. Total microbial DNA was isolated using the FastDNA® SPIN Kit (MP Biomedicals, USA) according to the manufacturer’s protocol. The quantity and quality of the DNA were checked, and DNA was then stored at −80 °C until use.

The community diversity was determined via sequencing 16S rRNA amplified with different primer pairs. (1) For bacteria, 338F (5’-ACTCCTACGGGAGGCAGCAG-3’) and 806R (5’-GGACTACHVGGGTWTCTAAT-3’) were used (Hamady et al., 2008). (2) For archaea, 524F-10-ext (5’-TGYCAGCCGCCGCGGTAA-3’) and 958R-mod (5’-YCCGGCGTTGAVTCCAATT-3’) were used (Pires et al., 2012). (3) For methanogens, MLfF (5’-GGTGGTGTMGGATTCACACARTAYGCWACAGC-3’) and MLrR (5’-TTCATTGCRTAGTTWGGRTAGTT-3’) were used (Luton et al., 2002). Purified amplicons were pooled in equimolar and paired-end sequenced (2 × 300) on an Illumina MiSeq platform (Illumina, San Diego, USA) according to the standard protocols by Majorbio Bio-Pharm Technology Co. Ltd. (Shanghai, China).

Sequence reads were assigned to samples by their nucleotide barcodes, merged according to their overlap by FLASH (v1.2.7, https://sourceforge.net/projects/flashpage/), and quality filtered (>220 bpwith less than 3% low-quality bases) by Trimmomatic-0.30 (http://www.usadellab.org/cms/index.php?page=trimmomatic). Operational taxonomic units (OTUs) were clustered with 97% similarity cutoff using UPARSE (version 7.1, http://drive5.com/uparse/) with a novel ‘greedy’ algorithm that performs chimera filtering and OTU clustering simultaneously. The taxonomy of each 16S rRNA gene sequence was analyzed by RDP Classifier algorithm (http://rdp.cme.msu.edu/) against the Silva (SSU123) 16S rRNA database using confidence threshold of 70%. LEfSe (Linear discriminant analysis effect size) analysis was performed by Galaxy modules of Huttenhower lab (https://huttenhower.sph.harvard.edu/galaxy/) with threshold LDA score of 4.0.

### Determination of *mcrA* levels by quantitative real-time PCR

The *mcrA* was quantitated by SYBR Green I-based qPCR method using the primer pairs, mlas (GGTGGTGTMGGDTTCACMCARTA) and mcrA-rev (CGTTCATBGCGTAGTTVGGRTAGT) (Steinberg and Regan, 2008). A 20 μL reaction contained 10 μL SYBR®Premix Ex Taq™ (Takara, China), 1 μL template DNA (5-10 ng), 0.4 μL (10 μM) of each primer, 0.4 μL of BSA (0.8 μg uL^-1^at the final concentration), 0.4 μL of ROX reference dye (50×) and 7.4 μl of sterile distilled water. A serials of known copy numbers of linearized plasmid DNA with the *mcrA* inserted from pure clones was used as standards for the quantitation. The thermal cycling was performed as following: 95 °C for 2 min, followed by 40 cycles of 95 °C for 5 s, 55 °C for 30 s, and 72 °C for 30 s. At least three biological repeats were prepared for each insect species, and three technical replicates were conducted for each qPCR reaction.

### Statistics

All data were analyzed with Data Processing System (DPS) software v9.50. The significance of differences was determined by one-way analysis of variance (ANOVA) and the significance level was set at 0.05 level.

## Data and materials availability

The raw reads were deposited into the NCBI Sequence Read Archive (SRA) database (Accession Number: PRJNA524105).

## Acknowledgements

We would like to thank Mrs Yan Lin (Nanjing Termite Control Institute, Nanjing, China) for the provision of *Coptotermes chaohuensis*. We would like to thank Professor Neil S. Millar (University College London, Gower Street, London, United Kingdom) for his help in improving the language. The work is supported by National Natural Science Foundation of China (grant number 31830075, 31772185 and 31701823).

