## Supplementary material for "Neonicotinoids stimulate H_2_-limited methane emission in *Periplaneta americana* through the regulation of gut bacterium community": Table S1, Table S2, Fig. S1, Fig. S2,Fig. S3, Fig. S4, Fig. S5, Fig. S6

**Supplementary tables**

**Table S1. Toxicities of cycloxaprid and imidacloprid against *Periplaneta americana***

| Insecticide | Abbreviation | *LC*_50_ (mg/L) | Slope |
| --- | --- | --- | --- |
| Cycloxaprid | CYC | 43.335 (35.725-52.555) | 2.813±0.242 |
| Imidacloprid | IMI | 57.030 (46.593-66.788) | 2.466±0.257 |

**Table S2. Toxicities of cycloxaprid and imidacloprid against *Coptotermes chaohuensis***

| Insecticide | Abbreviation | *LC*_50_ (mg/L) | Slope |
| --- | --- | --- | --- |
| Cycloxaprid | CYC | 5.136 (4.446-6.042) | 1.967±0.224 |
| Imidacloprid | IMI | 4.772 (4.215-5.536) | 2.124±0.195 |

**Supplementary figures**

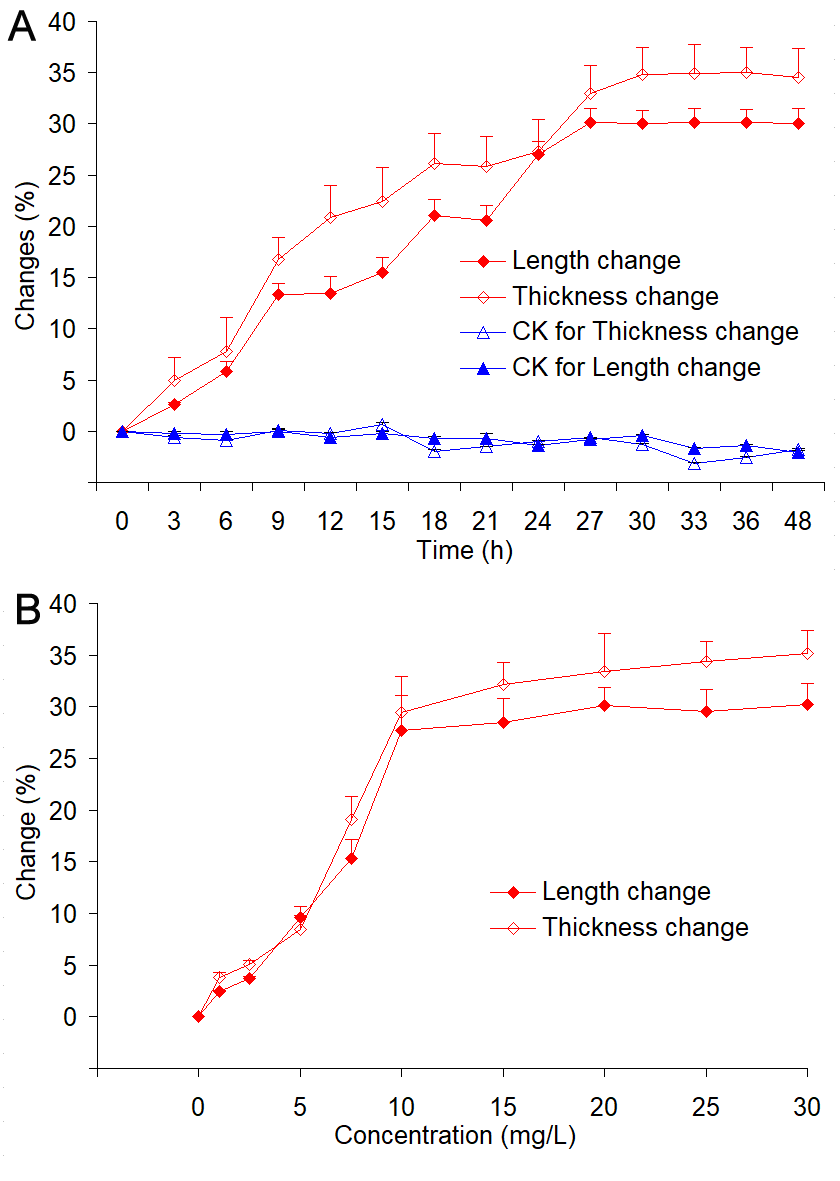

**Figure S1. The time- and concentration-dependent changes in the body length and thickness of *Periplaneta americana* treated by cycloxaprid (CYC).** (A) Changes in body length and thickness of *Periplaneta americana* at different time points after CYC treatment at *LC*_20_ concentration. (B) Changes in body length and thickness of *Periplaneta americana* treated by different CYC concentrations. Data are mean±SEM from at least five repetitions.

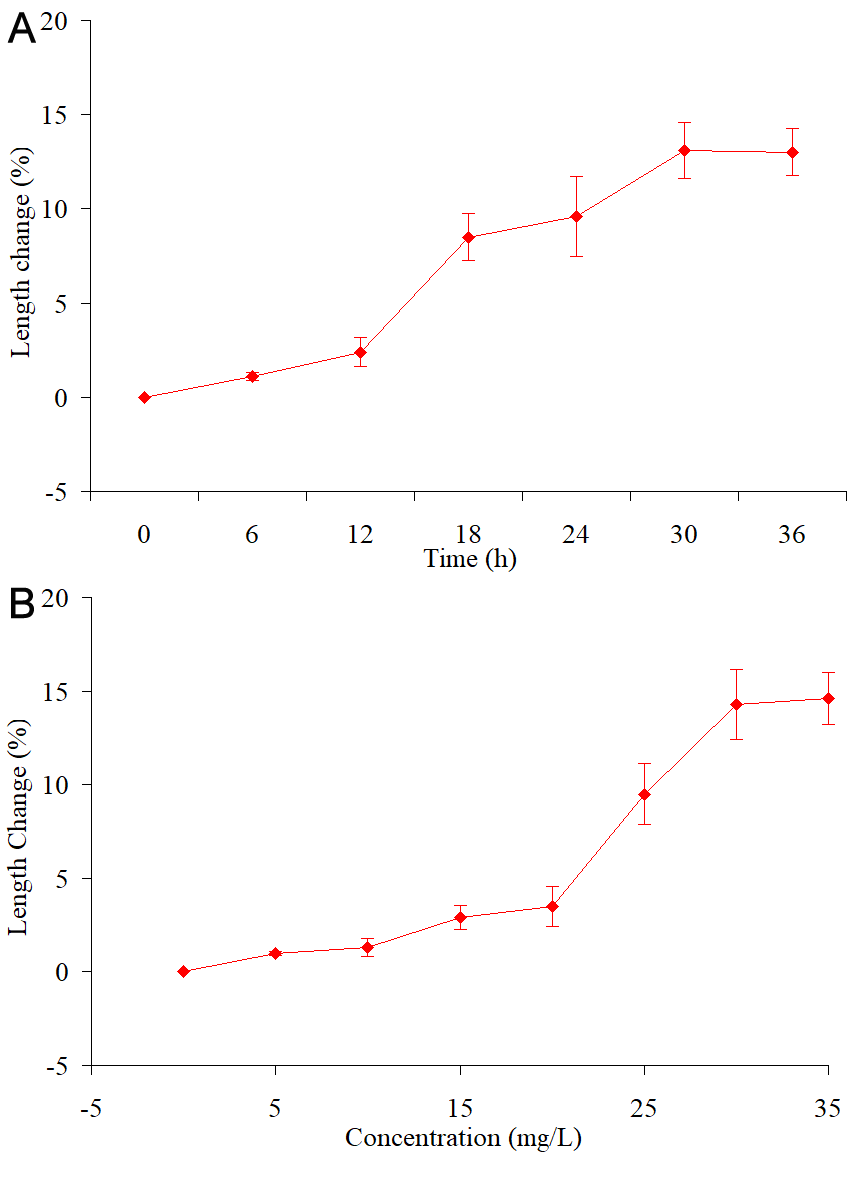

**Figure S2. The time- and concentration-dependent changes in the body length of *Periplaneta americana* treated by imidacloprid (IMI).** (A) Changes in body length and thickness of *Periplaneta americana* at different time points after IMI treatment at *LC*_20_ concentration. (B) Changes in body length of *Periplaneta americana* treated by different IMI concentrations. Data are mean±SEM from at least five repetitions.

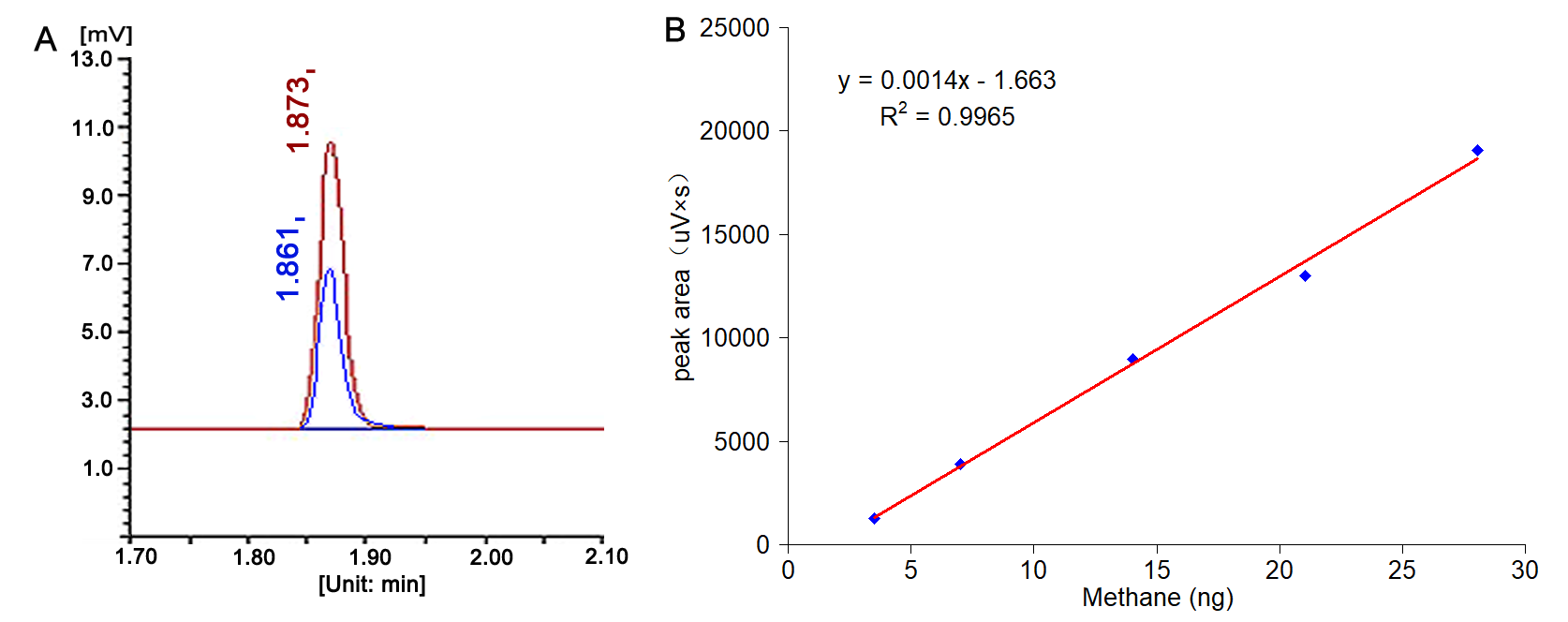

**Figure S3. Detection of methane and construction of the calculation equation between methane mass and areas in chromatographic recording.** (A) The gas chromatography identification of methane in the bloated foreguts of *P. americana* following cycloxaprid treatments. The red line marks methane from foreguts, and blue line marks the standard methane.

(B) The calculation equation between methane mass and chromatography areas. In the equation, y is methane mass (ng) and x is peak area (uV×s).

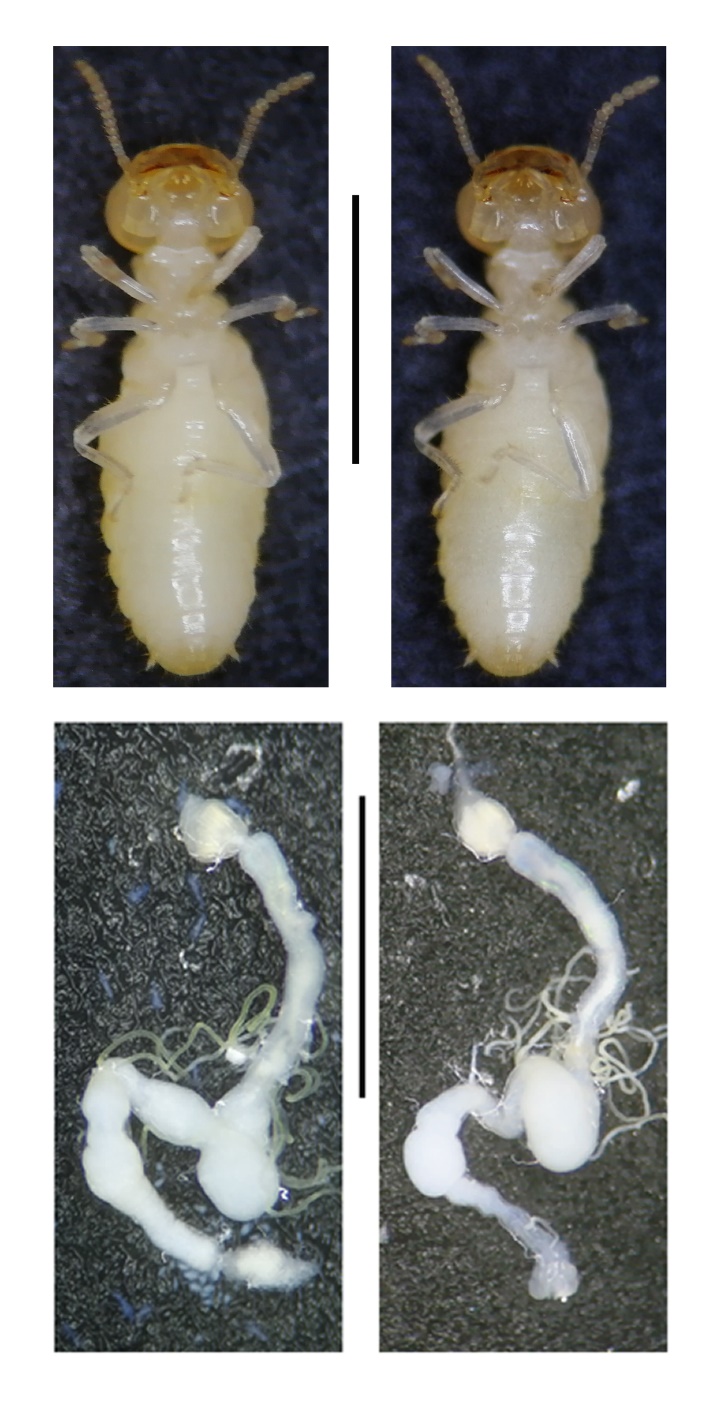

**Figure S4. The comparison of body shape and dissected guts of *Coptotermes chaohuensis* between the cycloxaprid treatment and untreated control.** The vertical line indicates a scale of 0.5 cm.

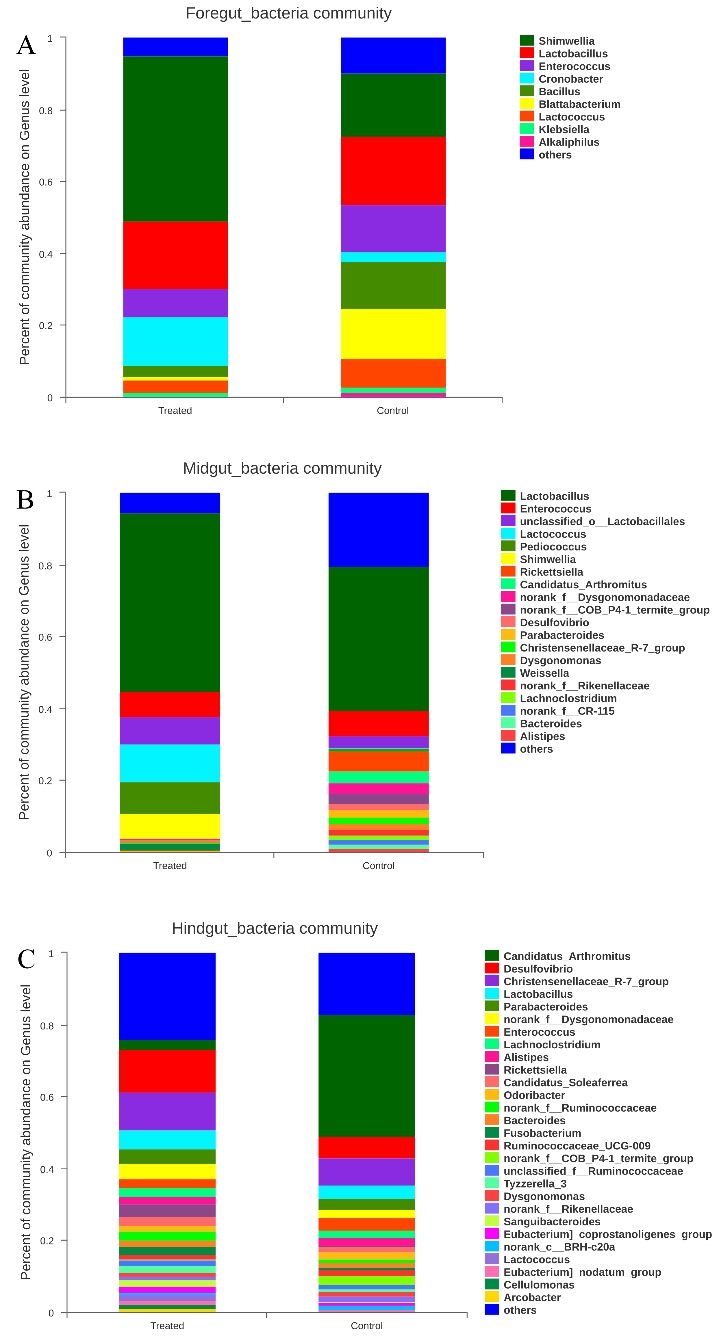

**Figure S5. The composition of bacterium community in three parts of *Periplaneta americana* guts.** (A) Foregut. (B) Midgut. (C) Hindgut. Data are average value of three independent samples.

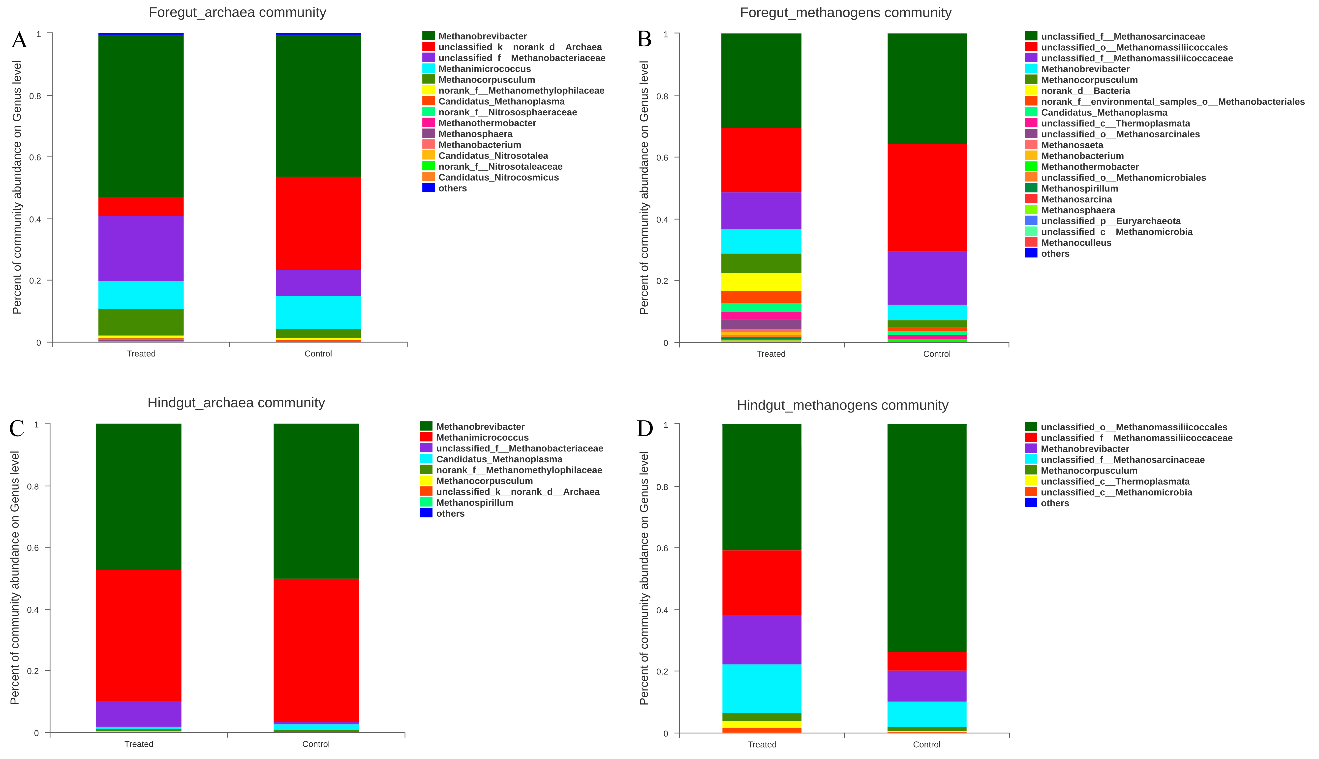

**Figure S6. The composition of archaea and methanogen communities** **in three parts of *Periplaneta americana* guts.** (A) Archaea in foregut. (B) Methanogens in foregut. (C) Archaea in hindgut. (D) Methanogens in hindgut. Data are average value of three independent samples.
